# Isoprenoid quinone profiling using a novel semi-quantitative HPLC-MS/MS method: application to wastewater sludges

**DOI:** 10.64898/2026.04.08.717208

**Authors:** Morgane Roger-Margueritat, Arthur Réveillard, Caroline Plazy, Andrei Octavian Filimon, Ahcène Boumendjel, Volker F. Wendisch, Valérie Cunin, Sophie S. Abby, Audrey Le Gouellec, Fabien Pierrel

## Abstract

Isoprenoid quinones are ubiquitous redox lipids that mediate electron transfer in various cellular processes across all domains of life. These molecules also serve as taxonomic and metabolic markers, facilitating the characterisation of microbial communities. However, their structural diversity and extreme hydrophobicity are challenging for comprehensive detection and quantification in complex biological matrices. In this study, we present a semi-quantitative HPLC-MS/MS method that enables the sensitive analysis of the widest range of quinones reported to date. Using a 16-quinone standard mixture, we optimized separation within a 14-minute HPLC gradient and achieved a sensitivity ranging from 55.2 to 0.3 femtomoles in targeted analyses. When applied to sewage sludges sampled weekly over three weeks, our method detected 57 quinones, revealing distinct quinone profiles at various wastewater treatment stages. This rapid and sensitive workflow enables quinone profiling in complex samples, and holds significant promise for advancing microbial ecology, environmental monitoring, and biotechnological applications.

## 1 INTRODUCTION

Isoprenoid quinones are a diverse family of redox-active lipids that are central to essential biological processes across all domains of life [1,2]. Isoprenoid quinones (hereafter referred to as « quinones ») feature a quinone ring conjugated to a polyprenyl chain whose length and saturation vary considerably among organisms, with some reaching up to 16 isoprene units [3,4]. The most prevalent quinones, such as ubiquinone (coenzyme Q) and menaquinone (vitamin K), are widely recognised for their roles in human health and nutrition [5–7].

Numerous methods have been described for quantifying selected quinones in various biological samples. Traditional methods involve extracting quinones by either liquid-liquid extraction of neutral lipids [8] or supercritical fluid extraction [9,10], followed by separation via capillary electrophoresis [11], gas chromatography [12] or, most commonly, high-performance liquid chromatography (HPLC) using C18 columns [13–16]. Detection techniques include UV absorbance, fluorescence, chemiluminescence, electrochemistry, mass spectrometry (MS) [16]. All these methods are limited to targeted analyses of a few quinones, such as coenzyme Q10 or vitamin K [17–20], and cannot capture the full diversity of quinones in complex microbial samples due to chromatographic overlap and constraints in sensitivity and selectivity.

Microbial quinone profiling has emerged as a powerful culture-independent tool for monitoring microbial ecology [21–23]. Quinone analysis provides insights into the structure and function of microbial communities, helping to optimise processes such as wastewater treatment [24–26]. Early HPLC-UV methods, pioneered by Hiraishi, could quantify up to 26 quinones in wastewater sludges and microbial mats. However, they required time-consuming sample fractionation to separate ubiquinones (UQs) and menaquinones (MKs) [27–29]. More recent HPLC-MS methods have expanded the range of MKs analysed, detecting MK4:4 to MK13:13 (i.e. MKs with chains containing 4 to 13 isoprene units) in human faeces with picomolar sensitivity [30–32]. However, these studies overlooked UQs and partially saturated MKs, which are produced by key bacterial genera [3]. The most advanced HPLC-MS/MS method can resolve 37 quinones in 42 minutes, including UQ7:7 to UQ10:10, MK6:6 to MK11:11 and several partially saturated MK [33]. A key strength of this method is the identification of quinones based on their fragmentation spectra, notably featuring the abundant tropylium ion characteristic of the quinone ring [33–35]. However, due to the limited commercial availability of quinone standards, reliance on a single standard to quantify the entire quinone series (MK4:4 for all MK and UQ10:10 for all UQ) was a critical limitation. This may introduce inaccuracies in the quantification given that MK isoprenologs ranging from MK4:4 to MK13:13 were reported to differ by up to tenfold in their MS response [30].

Here, we present an HPLC-Orbitrap method that sets a new standard for quinone profiling. Our chromatographic method only lasts 14 minutes and the combination of parallel reaction monitoring, high resolution masses and tropylium ion confirmation enabled the tentative annotation of 89 quinones. Quantification was possible for 13 authentic quinone standards (either commercially sourced or purified from microbial cultures), while semi-quantification was achieved for the other quinones. When applied to wastewater sludges, our method detected 57 distinct quinones, revealing significant diversity across treatment stages. Our HPLC-Orbitrap method overcomes some existing limitations and opens up new possibilities in microbial ecology and environmental monitoring by enabling the quantitative and semi-quantitative analysis of the most extensive quinone profile to date.

## 2 MATERIAL and METHODS

### 2.1 Quinone standard mixture

#### 2.1.1 Commercial quinones

UQ4:4, UQ10:10, MK7:7 were purchased from Sigma, UQ2:2 and UQ8:8 were obtained from LGC Standards, and MK9:9 was from Roth. MK4:4 and PK were from Tokyo Chemical Industry. Deuterated UQ10:10 (UQ10:10-d9, dimethoxy-d6 methyl-d3) was from Cambridge Isotope Laboratories, and MK1:1-d7, MK4:4-d7, MK9:9-d7 were from Toronto Research Chemicals. The quinones were suspended in hexane (at about 1 mM) and stored at -80°C, without any alteration for at least two years. The concentration of each solution was determined by UV-vis spectroscopy (Eppendorf BioSpectrometer) after dilution in ethanol for UQs or in hexane for MKs and PK. A molar extinction coefficient of 18800 cm^-1^ M^-1^ at 248 nm [36] was used to quantify MK and PK solutions, while molar extinction coefficients of 13700-15200 cm^-1^ M^-1^ at 275 nm were used for UQs [37]. We note that the use of generic molar extinction coefficient for MKs of various lengths will only cause a marginal uncertainty over the MK4-MK12 series, as response coefficients varied by less than 3% [27]. Solutions of each quinone were prepared at 200 µM in ethanol, and were then used to obtain an equimolar mixture at 10 µM in ethanol. Ten-fold serial dilutions in ethanol yielded the solutions at 1 µM, 100 nM, 10 nM, 1 nM and 0.1 nM which were used to generate the standard curves. XLogP3 values were obtained from Pubchem (https://pubchem.ncbi.nlm.nih.gov/).

#### 2.1.2 Purification of UQ6:6 from *Saccharomyces cerevisiae*

42 g of fresh baker’s yeast (brand « l’hirondelle » bought at local store) were homogenised in 15 mL of 0.15 M KCl. The cells were lysed using two cycles of French press at 1500 psi. The cell suspension was divided between ten 50 mL falcon tubes, to which 1.5 mL of glass beads (0.5 mm diameter) and 20 mL methanol were added. The tubes were vortexed for 1 min, after which 17 mL of petroleum ether (40-60° boiling range) was added to each tube. Vortexing was repeated for 30 seconds and phase separation was achieved via centrifugation at 1000 g for 3 min at room temperature. The top phase from each tube was transferred to a 500 mL round-bottom flask. Quinone extraction was repeated by adding 17 mL petroleum ether to the remaining methanol phases. The second petroleum ether phase was combined with the first, and the extract was evaporated in a rotary evaporator at 40°C. The extract was then suspended in 3 mL of cyclohexane. Analysis of an aliquot by high performance liquid chromatography-UV-electrochemical detection-mass spectrometry (HPLC-UV-ECD-MS) confirmed that UQ6:6 was the only quinone and that ergosterol was abundant (Figure S1A). HPLC-UV-ECD-MS analysis was performed as described previously [38] using a C18 column (Betabasic-18, 5 µm, 4.6 × 150 mm, Thermo Fisher) at a flow rate of 1 ml/min, with the mobile phase consisting of 50% methanol, 40 % ethanol, and 10 % of a mixture of 90% isopropanol, 10% ammonium acetate (1 M) and 0.1% trifluoroacetic acid.

To separate UQ6:6 from ergosterol, silica column chromatography was employed. A prepacked silica chromatography column (Advion Interchim SiStdF0004) was set on an Interchim PuriFlash 4250 device and activated using 30 mL of cyclohexane at 5 mL/min. Half of the quinone extract was then deposited onto the silica column. Elution was performed at a flow rate of 5 mL/min using a cyclohexane:ethyl acetate gradient. This started with 1 min at 100:0, followed by a 20 min gradient from 100:0 to 90:10, and a final 20 min gradient from 90:10 to 70:30. The fractions were collected in test tubes and analysed by HPLC-UV-ECD-MS to identify those containing UQ6:6 (eluting at 12 min) or ergosterol (eluting at 20 min). The fractions containing pure UQ6:6 were pooled, evaporated under reduced pressure and the purified UQ6:6 was resuspended in hexane. The UQ6:6 concentration was determined using UV-visible spectroscopy (Eppendorf BioSpectrometer) and a molar extinction coefficient of 14700 cm^-1^ M^-1^ at 275 nm [37]. Approximately 500 µL of UQ6:6 at a concentration of 1.44 mM was obtained.

#### 2.1.3 Purification of long-chain MKs from *Corynebacterium glutamicum*

*Corynebacterium glutamicum* Ubi5-Pd cells were grown in media containing 25 µg/mL kanamycin, 5 µg/mL tetracycline, and 100 µg/mL spectinomicin to maintain plasmid selection [39]. First, a glycerol stock was spread onto a LB plate and incubated overnight at 30°C. Then, the cells were inoculated into two baffled shake flasks containing 50 mL of LB medium supplemented with 10 g/L glucose, and grown overnight. Subsequently, two baffled shake flasks containing 500 mL of CgXII liquid medium supplemented with 40 g/L glucose and 1 mM IPTG were inoculated at an OD600 of 1 and cultured for 48h at 30°C with shaking at 120 rpm. The cells were then collected by centrifugation for 1h at 4000 g and 4°C. The pellet (42.6 g of wet cell mass) was frozen at -20°C and stored until further use. The pellet was then resuspended in 10 mL of 0.2 M KCl by vortexing, after which the cells were lysed using two cycles of French press at 1500 psi. 3 mL aliquots were transferred into 50 mL heavy-duty glass tubes with screw caps (Kimble Kimax). 3 mL glass beads (0.5 mm diameter) were added to each tube, along with 8 mL of methanol per gram of cells. The tubes were vortexed twice for 1 min, with a 5 min sonication step in between (ultrasonic bath XUBA1, Grant). Petroleum ether (40-60° boiling range) was added to the cell suspension (6 mL per gram of cells), after which the contents of the tubes were homogenised by vortexing for 1 min and sonicating for 3 min. Phase separation was achieved via centrifugation at 1,740 x g for 5 min at room temperature. The top phase (petroleum ether) from each tube was transferred to a pre-weighed 500 mL piriform flask (Duran) and protected from light using aluminium foil. Quinone extraction was repeated by adding 17 mL of petroleum ether to the remaining methanol phases, vortexing, sonicating and centrifuging. The second and first petroleum ether phases were combined and evaporated in a rotary evaporator at 40°C. This yielded 214.9 mg of oil, which was stored overnight at 4°C.

A prepacked C18 chromatography column (Advion Interchim C18HP-F0004 5 mL) was set on an Interchim PuriFlash 4250 device, and activated using 20 mL of a methanol - water (20:80) solution at 5 mL/min. Half of the quinone extract (suspended in approximately 1.5 mL of a 90:10 methanol-chloroform solution) was deposited onto the C18 column. Elution was performed at a flow rate of 5 mL/min using a methanol:isopropanol gradient, starting with a 10 min gradient from 100:0 to 64:36, followed by 60 min at 64:36. The fractions were collected in test tubes protected from light with aluminium foil and were analysed by HPLC-ECD-MS to identify those containing quinones. This analysis was performed using a C18 column (Betabasic-18, 5 µm, 4.6 × 150 mm, Thermofisher) at a flow rate of 0.75 ml/min with the mobile phase consisting of 50% isopropanol, 10 % ethanol, 30% methanol and 10 % of a mixture of 90% isopropanol and 10% ammonium acetate (1 M) with 0.1% formic acid. The quinones eluted from the C18 Puriflash column in the following order, with some overlap: MK10:10, MK10:9, MK11:11 and MK11:10. The fractions containing the purest quinones were pooled in 100 mL round-bottom flasks (MK10:10 + MK10:9; MK11:10), while those containing quinone mixtures were pooled and purified again using the same setup. After rotary evaporation, the purified quinones were resuspended in hexane, and the concentration of each quinone was determined by UV-visible spectroscopy (Eppendorf BioSpectrometer), using a molar extinction coefficient of 18800 cm^-1^ M^-1^ at 248 nm [36], and a normalisation to the peak area of each quinone, as obtained by HPLC-ECD-MS (Figure S1B). We obtained 1.1 mL of the 0.572 mM MK11:10 standard solution which also contained 0.038 mM MK11:11, as well as 2 mL of the purified MK10:10 + MK10:9 solution (0.172 mM and 0.180 mM, respectively) which also contained 0.006 mM MK11:11.

### 2.2 HPLC-MS/MS method

#### 2.2.1 Chromatography method

The UHPLC used was a Vanquish Flex (ThermoFisher Scientific) equipped with a binary pump. Quinones were separated using a C18 column (Uptisphere Strategy C18-3, 3 µm, 75×2.1 mm; Interchim), coupled to a pre-column (Uptisphere HP RP, 3µm, 5×2.1mm; Interchim), at 40°C with a flow rate of 0.5 mL/min. The elution was performed using two mobile phases: A (isopropanol (IP) HPLC-MS grade, containing 0.1% formic acid and 5mM ammonium acetate) and B (methanol HPLC-MS grade, containing 0.2% formic acid and 5mM ammonium acetate). The ammonium acetate stock solution used to prepare mobile phases A and B was 1M in ultra-pure water. The chromatography method was as follows. First, an isocratic step was carried out with 100% of phase B for 1 min, followed by a linear gradient from 100% to 25% phase B over 11 min. Then, phase B was decreased from 25% to 18% over 0.1 min, followed by an isocratic step at 18% B for 4.4 min. Finally, the column was equilibrated back to the initial condition by increasing B from 18% to 100% over 0.1 min, followed by an isocratic step at 100% B for 3.4 min. The total run time was 20 min.

#### 2.2.2 MS/MS method

A Q Exactive Plus Orbitrap high-resolution mass spectrometer (Thermo Fisher Scientific, Waltham, MA, USA) was employed with a heated electrospray ionisation (HESI) source operating in positive mode. Direct infusion of UQ10:10 and MK7:7 was first used to optimize parameters for efficient detection and fragmentation. The sheath gas flow rate was set to 60 and the auxiliary gas flow rate to 12. The spray voltage was set to 4.5 kV and the S-lens RF level to 70. The source temperature was set to 275°C for the capillary and 100°C for the auxiliary gas heater.

The untargeted Full MS / dd-MS² (TopN) method fragmented the 10 most abundant features (N = 10) simultaneously. A window covering the entire elution time was set for positive mode. For the full MS module, the resolution was set to 70,000 and the AGC target to 1e6. The maximum injection time was set to 150 ms, with a scan range of 200-1500 *m/z*. For the dd-MS² module, the resolution was set to 17,500, with multiple (N)CE at 10 V, 30 V and 50 V. The maximum injection time was set to 200 ms, and the isolation window to 1.5 *m/z*. The AGC target was configured for positive mode at 1e5. For the dd settings, the minimum AGC target was set to 1e4, the apex trigger was set between 0.1 and 55 s and the dynamic exclusion was configured to 5 s. Isotope exclusion was enabled.

In the targeted PRM method, each compound was searched according to an inclusion list (Table S1). A PRM window was set for positive mode throughout the elution time. The resolution was set at 17,500 with an (N)CE of 30 V. The maximum injection time was set to 100 ms, with an isolation window of 0.5 *m/z*. The AGC target was configured for positive mode at 1e5 and for negative mode at 5e4.

### 2.3 Sample collection and preparation of sludge samples

Samples were collected in the wastewater treatment plant of Romans-sur-Isère, France, once per week in June 2025, over a period of 3 weeks. Primary and activated sludges were sampled in their specific tanks in triplicate, below the surface (∼1m depth), while dehydrated (duplicate) and thickened (triplicate) were obtained from the centrifuges and the thickeners respectively, via dedicated collection pipes.

All sludge samples were put at 4°C for temporary storage in sealed polyethylene bottles, then transferred at -20°C, with aliquots to be analysed kept at -80°C. Sewage samples were centrifuged in 1.5 mL Eppendorf Safe-Lock tubes at 15,200 x g, 10 min, 4°C and 10 mg pellets (wet weight) were used for quinone extraction. The following was added to each 10 mg pellet: 100 µL of glass beads (0.5 mm diameter), 50 µL of 0.15 M KCl, 5 µL of a 20 µM solution of deuterated quinones (MK4:4-d7, MK9:9-d7 and UQ10:10-d9 diluted in HPLC grade ethanol), and 350 µL of distilled water. The samples were vortexed for 7 min at 1850 rpm (Digital Cell Disruptor, Scientific Industries), after which 600 µL of a mixture of isopropanol and hexane (ratio 3:2) was added. The samples were vortexed again for 3 min and sonicated for 2 min (ultrasonic bath XUBA1, Grant). The phases were separated by centrifugation for 1 min at 15,200 x g. The upper phase was transferred to a fresh tube. Then, 600 µL of the isopropanol:hexane mixture (ratio 3:2) was added to the remaining samples, and the extraction was repeated with vortexing, sonicating and centrifugating the samples. The upper phase was collected and combined with the previous one. A third extraction was then performed by adding 300 µL HPLC-MS grade hexane, vortexing for 2 min and centrifuging for 1 min. The upper layers were combined and dried under nitrogen. The dried extracts were resuspended in 100 µL of HPLC-grade ethanol and 5 µL were injected for analysis.

To determine the relationship between wet and dry weight for the different sludge types, five 10 mg aliquots of each type of sludge were transferred to pre-weighed 1.5 ml Eppendorf Safe-Lock tubes. The tubes were placed in an oven at 90°C for 2 hours with the lids open, followed by a further 2 hours in a drying incubator (Memmert BE 300) at 60°C. Empty control tubes (C1–C3) were treated similarly. The dry weights were measured after 30 minutes at room temperature (Table S2).

### 2.4 Data acquisition

The samples were stored at 15°C in the autosampler and an injection wash of 20 µL/s of HPLC grade ethanol was set for 30 s between each sample injection. The volumes of standards injected were 2, 5 and 10 µL, and the samples were 5 µL, which correspond to the extracts of 0.5 mg of sludges. The reproducibility between days was excellent, with no significant variation observed between the analysis of standards or sludge samples up to three days apart.

Quality controls (QC) were performed to check the quality of the acquisition. A QC-pool was created by pooling 5 µL of each sludge sample, and diluting it serially in ethanol (1/2, 1/4, 1/8 and 1/16 dilutions). 10 µL of each were injected every 10 samples to assess the linearity of the MS signals for all quinones. The five dilutions followed a linear regression, with a median CV of less than 15%. 10 µL of HPLC-grade ethanol were injected as blank samples after the lowest dilution of each QC series. No quinones were detected in the blank samples, demonstrating the absence of carry-over.

### 2.5 Data analysis

#### 2.5.1 Analysis of untargeted MS/MS data

The raw untargeted MS/MS data was converted to mzML files and processed using the MZmine software (v.4.9.14) with relative mass tolerance set at 5 ppm. Noise thresholds of 2.5e4 and 1.0e3 were set for MS1 and MS2 mass detections, respectively. Extracted ion chromatograms (EIC) were built using the chromatogram builder tool, with a minimum peak height intensity of 1.0e4 and chromatographic deconvolution (separation of detected masses into individual peaks) was carried out using the local minimum feature resolver algorithm where the minimum peak height was set at 5.0e4 and the maximum peak width at 2 min. Following the deconvolution, ^13^C isotopes were filtered with a 0.1 min retention time tolerance followed by alignment of EICs with an 8 ppm mass tolerance and a 0.3 min retention time tolerance. After correlation grouping (metaCorrelate) with a retention time tolerance of 0.1 min and a minimum feature height of 3.0e5, adducts identities were determined using the ion identity molecular networking (IIMN) module of MZmine, selecting [M+H]^+^ and [M+NH_4_]^+^adducts. Spectral annotation from reference libraries was eventually performed using publicly available libraries (MoNA, Mass BankEU, GNPS), as well as an in-house quinone-specific library. For the spectral library search, a minimum of 5 matched signals had to be detected with a minimum modified cosine similarity of 0.7 for a spectral match to be confirmed. A post-treatment filtering was applied with a maximum delta retention time of 0.15 min and a maximum delta *m/z* of 5 mmu (milli mass unit).

#### 2.5.2 Analysis of PRM data

The PRM raw data were analysed using the General Quan method in TraceFinder 4.1 (Thermo Fisher Scientific, Waltham, MA, USA). A master method was designed to confirm the presence of each compound in raw data files based on the detection of the adduct mass and the corresponding tropylium ion. The smoothing curve was calibrated at the minimum (1) and the detection algorithm was ICIS. The master method was then executed on all PRM raw data files, after which the output table was extracted. Manual curation was carried out to ensure the entire peak was integrated when it had not been fully recovered automatically.

A homemade script designed in Python (v.3.9.16) was then used to apply two cut-off values to the obtained TraceFinder output table: a maximum delta retention time of 0.15 min and a maximum delta *m/z* of 2 mmu. This step eliminated some features which were originally present in the TraceFinder output table, such as several DMKs and short chain MKs.

#### 2.5.3 Calculation of MS parameters

Following post-analysis treatment, the linearity of the signal of each quinone standard in untargeted and targeted was assessed and the matplotlib (v.3.5.1 [40]) module was used to plot data and the NumPy (v.1.26.4 [41]) module to compute the trendline and the linear correlation coefficient R². The ULOQ was defined as the highest quantity injected following the linear regression. The noise (N) was obtained from the untargeted and targeted data by retrieving the maximum noise value at the corresponding retention time for each standard from mzMine or Qual respectively. The LOD and LOQ were calculated as 3 × N and 10 × N, respectively. For the CV, the standard deviation of replicates at a given concentration was divided by the average obtained for these same replicates. The CV was calculated at all concentrations based on the LOD of each compound for PRM data or untargeted MS/MS data (n=3). The median of all CVs was then obtained. All these definitions are in accordance with the COFRAC guidelines.

#### 2.5.4 Quantification and semi-quantification of quinones

Quantitative values in pmoles per mg were obtained for sludge samples by correcting values for extraction loss based on the internal deuterated standards, by normalizing to sample weight, and by using a trendline coefficient adapted to each quinone (Table 1 and Table S3). For non-standard quinones, trendline coefficients used were obtained for MK and UQ by averaging the values of the two nearest quinone standards (Figure S7), except for quinones longer than MK11:11 and UQ10:10, for which the latter’s coefficient was used. Coefficients for the other types of quinones were obtained based on the structurally closest standard: UQ for mPQ, PQ, RQ and MQ; MK for PK, MMK and DMK.

**Table 1:** Targeted detection of quinones in the standard mixture. The parameters were computed for the most abundant adduct of each quinone, following the injection of the standard mixture in triplicate over a range of 0.0002 to 100 pmoles, except for MK11:11, which was approximately ten times less abundant; see Text S1 for details. This made it impossible to evaluate the ULOQ for MK11:11 above 10 pmoles.

| Compound | Exact mass | Ionized mass (m/z) | Adduct | Ionization mode | RT (min) | CV (%) | LOD (pmoles) | LOD (pg) | LOQ (pmoles) | ULOQ (pmoles) | Trendline coefficient | R<sup>2</sup> |
| --- | --- | --- | --- | --- | --- | --- | --- | --- | --- | --- | --- | --- |
| MK4:4 | 444.30283 | 445.3106 | H<sup>+</sup> | pos | 1.85 | 8.11 | 0.0408 | 18.11 | 0.1358 | 50 | 5.09E+06 | 0.988 |
| MK4:4 d | 451.34693 | 452.3548 | H<sup>+</sup> | pos | 1.85 | 6.31 | 0.0552 | 24.90 | 0.1839 | 50 | 6.02E+06 | 0.986 |
| MK7:7 | 648.49063 | 666.525 | NH4<sup>+</sup> | pos | 5.7 | 4.64 | 0.0033 | 2.14 | 0.0110 | 50 | 1.04E+07 | 0.997 |
| MK9:9 | 784.61583 | 802.6502 | NH4<sup>+</sup> | pos | 8.4 | 3.44 | 0.0023 | 1.77 | 0.0075 | 50 | 9.42E+06 | 0.995 |
| MK9:9 d | 791.65993 | 809.6943 | NH4<sup>+</sup> | pos | 8.4 | 3.69 | 0.0026 | 2.08 | 0.0088 | 50 | 1.36E+07 | 0.993 |
| MK10:10 | 852.67843 | 870.7128 | NH4<sup>+</sup> | pos | 9.55 | 3.47 | 0.0015 | 1.31 | 0.0051 | 46 | 1.05E+07 | 0.994 |
| MK10:9 | 854.69408 | 872.7284 | NH4<sup>+</sup> | pos | 10 | 3.24 | 0.0014 | 1.16 | 0.0045 | 50 | 1.23E+07 | 0.991 |
| MK11:11 | 920.74103 | 938.7754 | NH4<sup>+</sup> | pos | 10.55 | 3.31 | 0.0010 | 0.96 | 0.0035 | 10 or more | 1.52E+07 | 0.992 |
| MK11:10 | 922.75668 | 940.791 | NH4<sup>+</sup> | pos | 10.95 | 2.95 | 0.0018 | 1.62 | 0.0058 | 50 | 1.55E+07 | 0.997 |
| PK | 450.34963 | 451.3574 | H<sup>+</sup> | pos | 3.2 | 6.32 | 0.0386 | 17.39 | 0.1287 | 50 | 2.03E+06 | 0.987 |
| UQ2:2 | 318.18311 | 319.1909 | H<sup>+</sup> | pos | 0.6 | 5.07 | 0.0072 | 2.28 | 0.0239 | 20 | 2.24E+07 | 0.993 |
| UQ4:4 | 454.30831 | 455.3161 | H<sup>+</sup> | pos | 1.1 | 5.49 | 0.0011 | 0.48 | 0.0035 | 50 | 3.44E+07 | 0.994 |
| UQ6:6 | 590.43351 | 608.4679 | NH4<sup>+</sup> | pos | 2.6 | 5.95 | 0.0082 | 4.82 | 0.0272 | 100 | 1.26E+07 | 0.999 |
| UQ8:8 | 726.55871 | 744.5931 | NH4<sup>+</sup> | pos | 5.15 | 4.80 | 0.0025 | 1.79 | 0.0082 | 50 | 2.37E+07 | 0.998 |
| UQ10:10 | 862.68391 | 880.7183 | NH4<sup>+</sup> | pos | 7.75 | 4.39 | 0.0004 | 0.35 | 0.0014 | 20 | 3.72E+07 | 0.999 |
| UQ10:10 d | 871.74061 | 889.7749 | NH4<sup>+</sup> | pos | 7.65 | 3.80 | 0.0003 | 0.30 | 0.0011 | 20 | 5.43E+07 | 0.999 |

PCA were computed and visualized with the ggbiplot (v. 0.6.2) and ggplot2 (v. 3.5.2) of R with the best quinone descriptors displayed by black arrows. The ellipses for each sludge type represent the 68% confidence ellipses, which were calculated by ggbiplot for each group based on the individuals’ coordinates on the first two principal components. The Hellinger transformation was then applied using the decostand function from the vegan (v. 2.7.1) package.

Differences in the overall quinone profiles among sludge types were assessed using permutational multivariate analysis of variance (PERMANOVA), implemented with the adonis2 function from the vegan package. Euclidean distances were calculated from the Hellinger-transformed quinone data, and statistical significance was assessed using 999 permutations. The homogeneity of multivariate dispersions was then assessed using the betadisper function and evaluated using the anova and the permutest functions. Pairwise differences between sludge types using the TukeyHSD function were obtained to see which type of sludge had heterogeneous distribution. An additional PERMANOVA test was performed to assess the cumulative effect of the sludge type and the sampling week (999 permutations), as well as the dispersion with betadisper followed by permutest.

## 3 RESULTS

### 3.1 Untargeted LC-MS/MS approach for method development

Natural quinones exhibit a remarkable range of hydrophobicity, with XLogP3 values spanning from 7.1 (MK3:3) to 29.4 (MK15:15) [42]. To develop a robust method for their comprehensive profiling, we first needed a diverse and representative set of standards. We assembled a mixture of 17 quinones by combining (i) commercially available quinones (MK4:4, MK7:7, MK9:9, phylloquinone (PK), UQ2:2, UQ4:4, UQ8:8, and UQ10:10), (ii) deuterated analogues (MK1:1-d7, MK4:4-d7, MK9:9-d7 and UQ10:10-d9), and (iii) quinones purified from *Saccharomyces cerevisiae* (UQ6:6) and from *Corynebacterium glutamicum* Ubi5-Pd cells [39] (MK10:10, MK10:9, MK11:11 and MK11:10; see Text S1 and Figure S1).

We developed an HPLC method using a C18 column and a methanol/isopropanol gradient to optimize the separation of the quinone standards. Overlaying MS scans of individual quinones obtained by untargeted full MS/data-dependent-MS² (MS/dd-MS²) revealed efficient separation within 11 minutes (Figure 1A). However, despite equal concentrations in the mixture (except for MK11:11), peak areas varied significantly, highlighting differences in ionization efficiency. For example, MK1:1-d7 was nearly undetectable and was therefore excluded from further analysis, and phylloquinone (PK or MK4:1) exhibited much lower instrument response than MK4:4 (Figure 1A). As PK and MK4:4 differ only by three saturations on their prenyl chain, this suggests that chain saturation impacts ionisation efficiency, though this was not observed for long-chain MKs.

**Figure 1:**
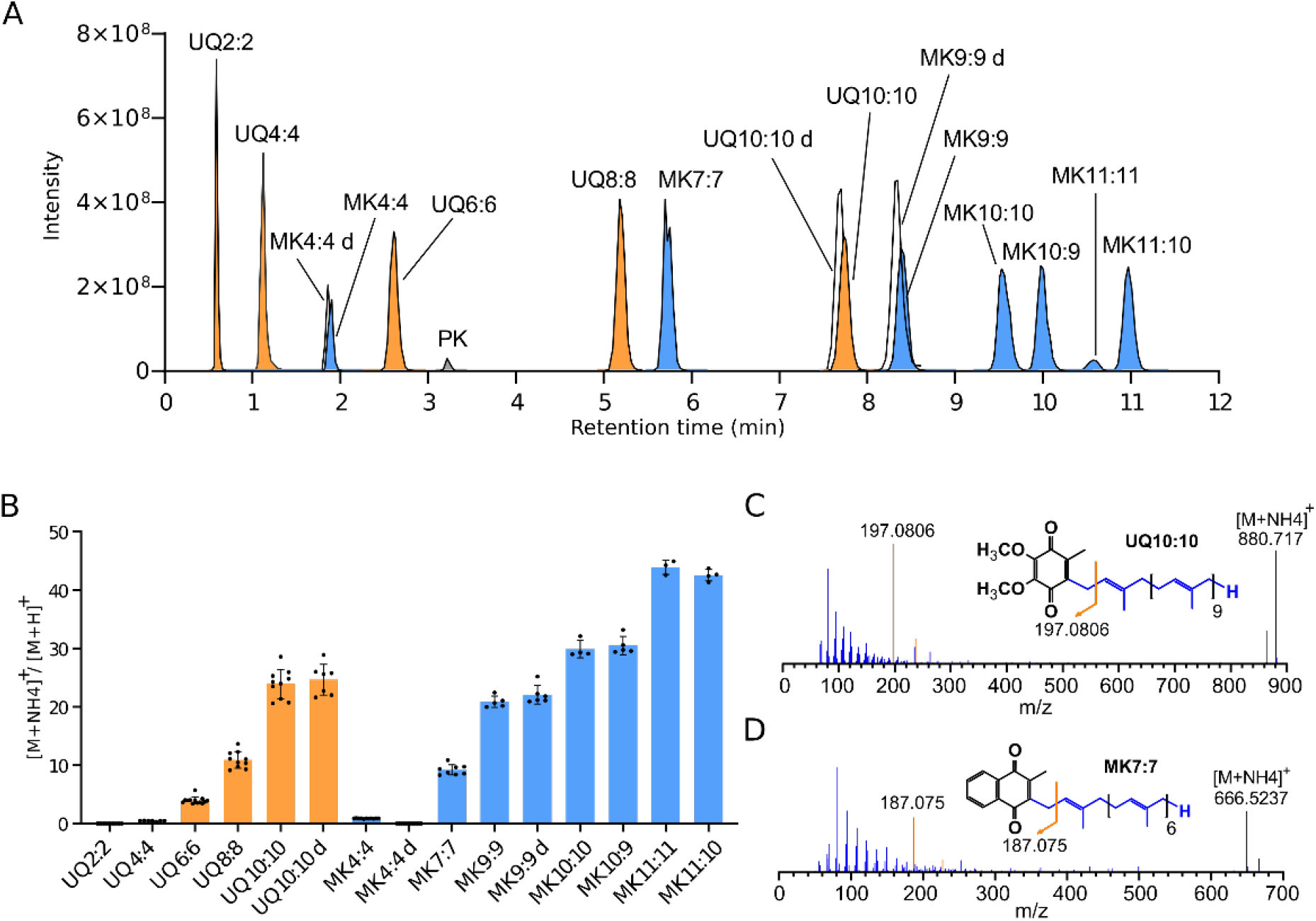
Quinone detection using an untargeted approach. **A**. Reconstructed chromatogram of detected ubiquinones (orange) and menaquinones (blue) after injection of 20 pmoles of the standard mixture. **B**. Ratio between NH_4_^+^ and H^+^ adducts for each ubiquinone (in orange) and menaquinone (in blue) from the standard mix. Mean ± SD (n = 4-6). C-D. MS/MS fragmentation spectra for ubiquinone (**C**, UQ10:10) and menaquinone (**D**, MK7:7), see also Figure S2. The tropylium ion peak is coloured in orange and the unfragmented quinone peak is coloured in black.

A notable finding was that adduct formation was influenced by chain length. Protonated species (H^+^) were dominant for shorter chains (up to UQ4:4 and MK4:4), whereas ammonium adducts (NH ^+^) prevailed for longer chains, with a difference up to 40-fold for MK11:11 and MK11:10 (Figure 1B). Interestingly, omitting ammonium acetate from the mobile phase did not increase the signals of protonated adducts. In line with previous reports [22,34,35], the fragmentation spectra displayed prominent tropylium ions at m/z 197.0806 (UQ) and 187.075 (MK) (Figures 1C-D and Figure S2).

To assess signal linearity, 2, 5 or 10 µL of the quinone standard mixture was injected at concentrations ranging from 0.1 nM to 10 µM (0.0002–100 pmoles injected). The signal areas obtained after MZmine processing were plotted against the injected amounts (Figure S3), selecting NH_4_ ^+^ adducts for quinones with five or more isoprene units and H^+^ adducts for quinones with shorter chains. The limit of detection (LOD), the limit of quantification (LOQ) and the upper limit of quantification (ULOQ) were calculated and are reported in Table S4. The results revealed that UQs were detected more efficiently than MKs (e.g. an LOD of 0.005 pmol for UQ10:10 versus 0.0184 pmol for MK10:10) and that long-chain quinones were detected more efficiently than short-chain ones (e.g. an LOD of 0.02 pmol for MK9:9 versus 0.05 pmol for MK4:4). Signal saturation occurred at high concentrations, leading to a ULOQ of 20 or 50 pmoles for most quinones (Table S4).

Despite compiling the most extensive collection of quinone standards to date, our set does not cover the full range of natural quinones [3,4]. In order to increase the diversity of quinones monitored by our method, we assembled a quinone spectral library based on untargeted analyses of various biological samples. In total, we identified 75 quinone-related features (Table S5, and spectral library available in Figshare). The level of confidence in the annotation of these compounds is high since the molecular mass of each compound differed from the theoretical mass by less than 3 ppm (and was often <1 ppm, Table S5), and characteristic fragments, such as the tropylium ion and several chain fragments were also observed with high mass accuracy (Table S5 and spectral library).

### 3.2 Development of a targeted LC-MS/MS method

To enhance sensitivity and enable the (semi)-quantification of trace quinones in complex matrices, we developed a targeted approach using parallel reaction monitoring (PRM), leveraging tropylium ions as diagnostic fragments. The PRM windows were designed to allow for a margin of at least 0.3 minutes on either side of the observed retention time (Figure S4, Table S1). The inclusion list comprised the 16 quinone standards, classified as level 1 according to the Metabolomics Standards Initiative (MSI) criteria [43], the 59 quinone features identified in the untargeted approach, classified as level 2, and some predicted quinones, such as MK8:4 or MK10:5, with polyprenyl chains different from those observed in the untargeted approach (Table S5). All those predicted quinone features were assigned an MSI level 2 (Table S5), since they were observed in subsequent PRM analyses, with retention times consistent with their chain length and saturation (Figure S4, Table S5).

Analysis of the standard mixture revealed detection patterns consistent with those observed in full MS/dd-MS² mode (Figures S5 and 1A). Raw PRM data were processed using TraceFinder 4.1, and peak areas were extracted for quantification (Figure S6). Stringent filters were applied to ensure robust identification: a retention time (RT) reproducibility window of 0.15 minute and a mass accuracy threshold of 2 millimass units. This approach effectively resolved potential interferences. For example, MK10:9 and MK11:10 could be distinguished from the ^13^C₂ isotopes of MK10:10 and MK11:11 based on their distinct RTs (differences > 0.15 min), despite their nearly identical masses.

The PRM method substantially improved the LOD and LOQ of the 16 quinone standards compared to the untargeted approach (e.g. 350 versus 1.1 fmoles for UQ4:4; see Table 1 and S1). Additionally, the ULOQ was extended for certain standards (Figure 2, Table 1), thereby broadening the dynamic range (e.g. 50 pmoles versus 20 pmoles for UQ4:4). All standards exhibited a median coefficient of variation (CV) below 10% (Table 1), which is well within the 15% threshold considered acceptable for quantitative analysis [33,34].

**Figure 2:**
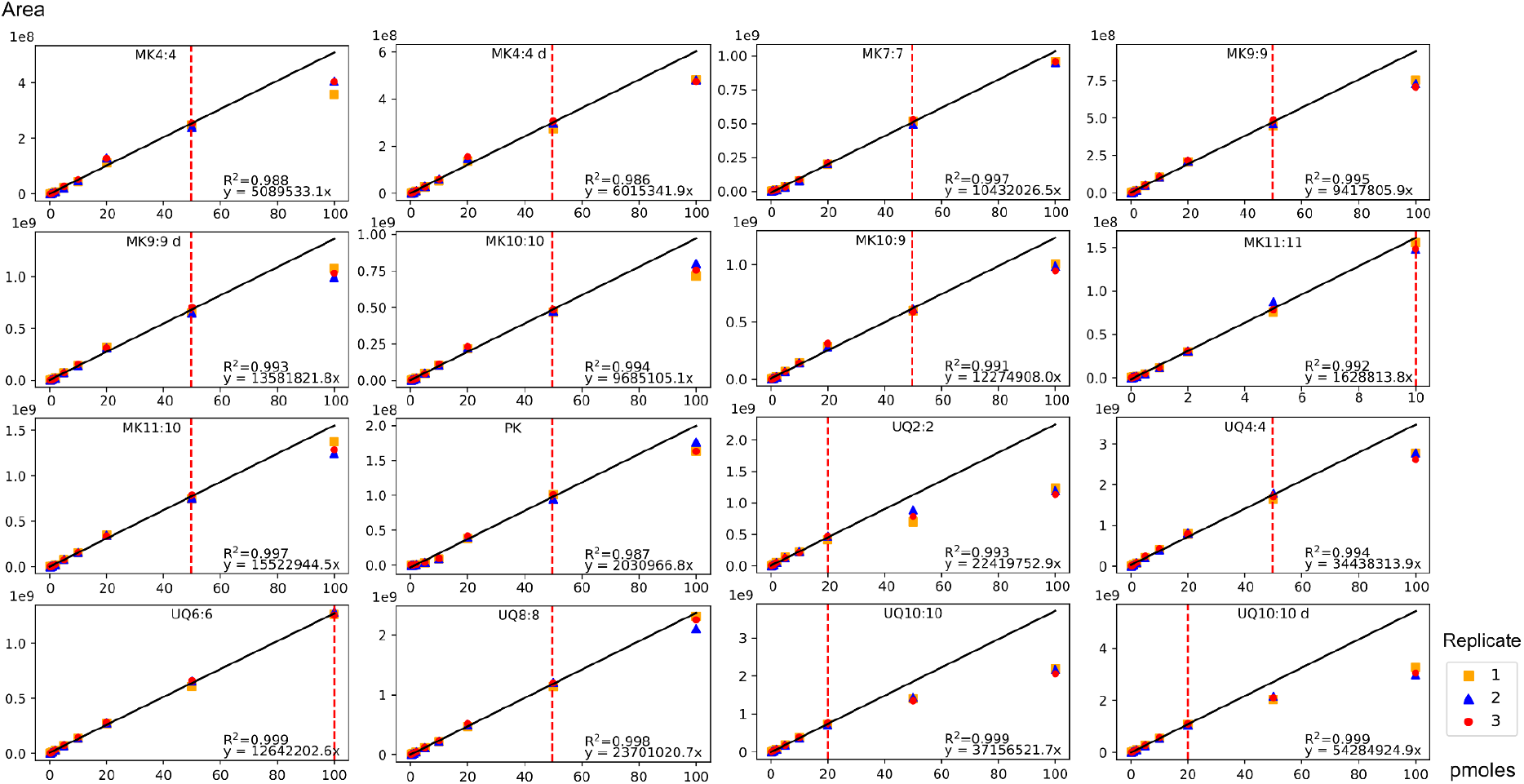
Linearity of quinone standards using targeted PRM approach. A mixture of quinone standards was injected over a range of 0.0002 to 100 pmoles in triplicates. The linear correlation coefficient (R²) and the corresponding trendline for the linear part of the curves were computed using the Python modules Numpy. The red dashed line corresponds to the Upper Limit of Quantification (ULOQ).

Notably, a linear correlation emerged between the trendline coefficient and chain length for the MK4:4–11:11 and UQ6:6–10:10 series of standards (Figure S7), with respective differences of 3.04- and 2.95-fold between the lowest and highest values. To enable the semi-quantitative analysis of quinones not available as standards, we estimated their response factor by averaging the values of the two nearest available quinone standards, or by applying the values of the structurally similar quinones. For example, we used the value of MK9:9 for DMK9:9 and of UQ9:9 for mPQ9:9 and PQ9:9 (see Figure S7, Methods and Table S3).

### 3.3 Quinone extraction from wastewater sludges and matrix effect

As wastewater sludges are known to contain a large variety of quinones [28], this type of sample was chosen to apply our PRM method. Four types of sewage sludge (primary, thickened, activated and dehydrated) were collected weekly from a neighbouring wastewater treatment facility over a period of three weeks.

To evaluate the efficiency of the extraction process and the impact of the matrix effect, which is here interference of the sludge matrix, deuterated quinones were spiked before or after the extraction step respectively. While extraction losses remained minimal for all three standards, a pronounced matrix effect was observed for MK4:4-d7, resulting in recovery rates as low as 14% (Figure S8). However, no such effect was observed for MK9:9-d7 or UQ10:10-d9, with recovery rates of 93.7% and 100% respectively (Figure S8). These results suggest that short-chain quinones are disproportionately impacted by matrix interference, whereas long-chain quinones can be more reliably detected and quantified in complex sewage matrices.

### 3.4 Application of the targeted LC-MS/MS method to wastewater sludges

Out of the 89 quinones targeted by our PRM approach, 57 quinones were detected across the 51 analysed samples, ranging UQ2:2-UQ12:12 and MK4:4-MK15:15 (Figure 3A, Table S6). Quantification was achieved for the quinone standards, but not for MK4:4, MK10:9, MK11:10, PK, UQ2:2 and UQ:4:4, as these were below the LOQ for some sludge samples. Semi-quantification was achieved using estimated response factors for the quinones for which there were no authentic standards (see Figure S7, Table S3, and Methods). In line with previous studies [25,27,28], the most abundant benzoquinones were UQ8:8 and UQ10:10, and the predominant naphthoquinones were MK7:7, MK8:8 and MK8:6 (Figure 3B).

**Figure 3:**
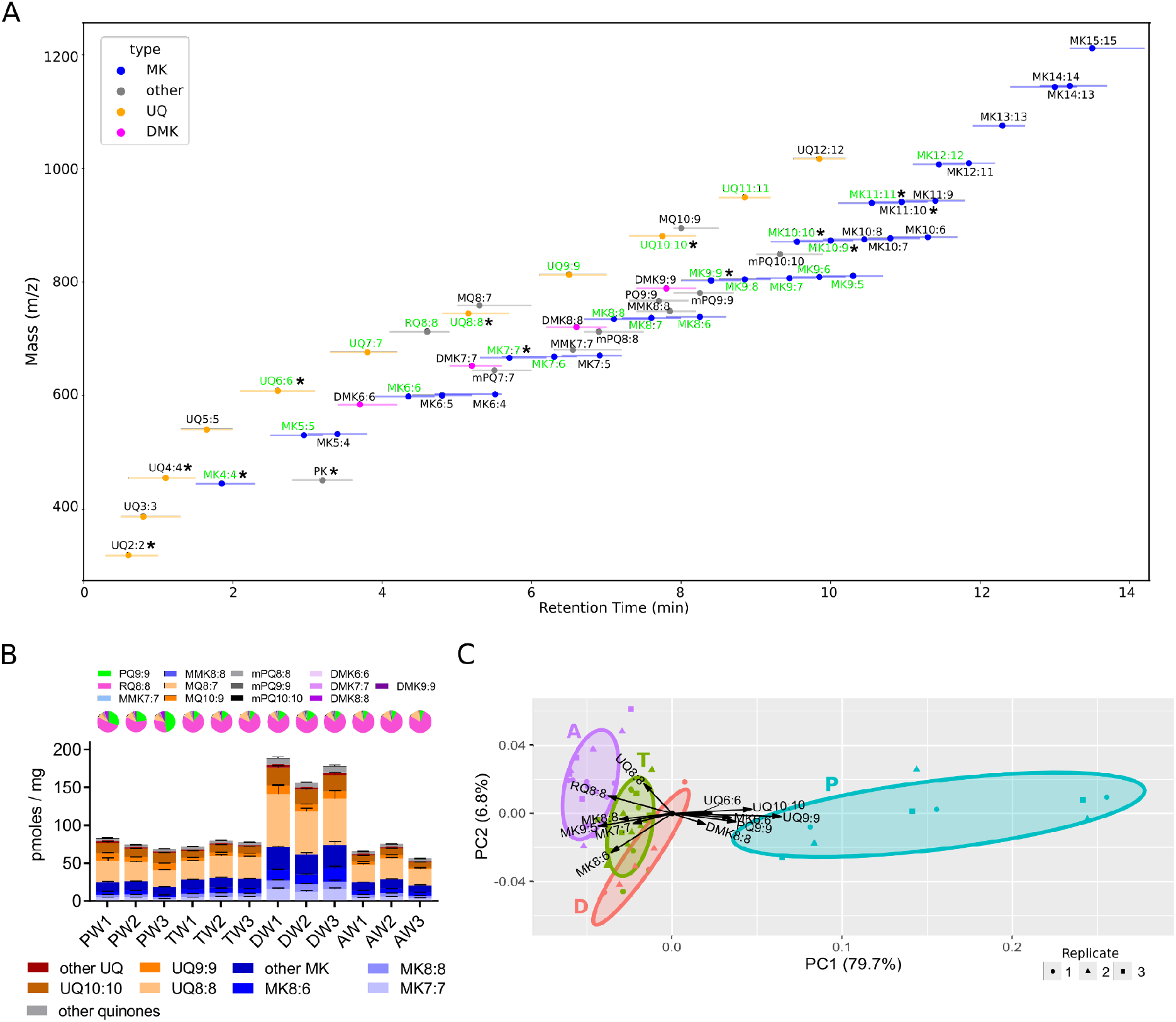
Quinone detection in wastewater sludges. **A**. PRM windows for quinone detected in sludge samples. The dots represent the average retention time of each quinone, and lines represent the span of the PRM windows. Quinones detected previously in sludges [25] are indicated in green. Confirmed quinones (standards) are annotated with a black asterisk whereas the others are putatively annotated according to their fragmentation pattern and their mass (Table S5). UQ, ubiquinone; DMK, demethylmenaquinone; MK, menaquinone; MMK, methylmenaquinone; PK, phylloquinone; RQ, rhodoquinone; mPQ, methyl plastoquinone; PQ, plastoquinone; MQ, methylene ubiquinone. **B**. Quantified and semi-quantified quinone profiles from primary (P), thickened (T), activated (A) or dehydrated (D) sludges retrieved over three consecutive weeks (W1, W2 or W3). Quinones representing less than 5% of the total across all samples were grouped as “other MK”, “other UQ” or “other quinones”. Values below the LOQ were not displayed and the remaining values are presented as mean + SEM (n = 2-6). The pie plots represent the “other quinones” category with the legend shown above. **C**. Principal component analysis (PCA) after Hellinger transformation of quinones detected in each sludge type: primary (P), thickened (T), activated (A) and dehydrated (D). See also Figures S9-S12.

The quinone profiles remained consistent throughout the three-week sampling period (Figures 3B and S9). Principal component analysis (PCA) distinguished dehydrated sludges from the other sample types based on the abundance of UQ8:8, UQ9:9 and UQ10:10 (Figure S10). To mitigate the influence of quinone abundance and account for variability in quinone types, Hellinger transformation was applied prior to PCA [44] (Table S6). This separated the samples according to their treatment stage, particularly the primary sludges (Figure 3C). Interestingly, sampling time contributed minimally to variability, except in primary sludges where weekly triplicates clustered along the first PCA component (Figure 3C, Figure S11). The quinone profiles differed significantly between sludge types (PERMANOVA controlling for the week effect: R² = 0.723, p = 0.001). Although the multivariate dispersions differed between sludge types (PERMDISP: p = 0.001), the effect of sludge type was considerably greater than the week effect (R² = 0.040), indicating that the observed pattern is primarily associated with the treatment process. A PCA focused on primary sludges revealed that week 3 samples were characterised by higher levels of UQ10:10, UQ9:9, PQ9:9, and MK6:6, whereas week 2 samples were distinguished by unsaturated MKs (Figure S12).

Together, these results reveal stage-specific quinone profiles in the wastewater treatment process. Primary sludges exhibited the greatest week-to-week variability, likely reflecting fluctuations in influent composition.

## 4 DISCUSSION

The HPLC-MS/MS method presented here is a significant advancement in quinone profiling. It enables the sensitive analysis of the widest range of quinones reported to date, from UQ2:2 to MK15:15, in just 14 minutes of chromatographic separation (20 minutes including column re-equilibration). This constitutes a twofold reduction in analysis time compared to the current state-of-the-art HPLC-MS/MS method [33]. Using a set of 16 purified quinone standards, we found that short-chain quinones predominantly form H^+^ adducts, whereas long-chain quinones are more likely to generate NH_4_ ^+^ adducts (Figure 1B). We note that this phenomenon may be influenced by factors other than quinone chain length, such as gradient composition, desolvation, and competition for ionisation. Detection sensitivity also depended on chain-length, with LOD values being approximately twice as high for short-chain than for long-chain MKs and UQs (Table 1). Similarly, the response factors varied by approximately threefold within the MK4:4–11:11 and UQ6:6–10:10 series of standards (Figure S7 and Table 1), indicating that using a single standard for quantification, as was sometimes done in previous analyses, is not satisfactory. Currently, the best way to achieve semi-quantitative analyses of quinones unavailable as standards is to estimate their response factors, despite this introducing significant uncertainty in the results. This limitation could be overcome in the future by obtaining a more comprehensive set of standards in order to enable fully quantitative analyses across the entire quinone spectrum. As several quinone types, including DMK, PQ and mPQ, are not commercially available, it is feasible, albeit time-consuming, to purify them from natural sources using microorganisms that can be cultivated on a large scale.

Another limitation lies in the matrix effect, which was particularly pronounced for short-chain quinones in sludge samples (see Figure S8). As it would be difficult to extrapolate the matrix effect for non-standard quinones, we decided not to adjust the values in the sludge samples dataset (Table S6). This may result in certain quinones being underestimated, particularly those with short chains. Consequently, this dataset should be considered primarily for its comparative value, *i.e.* the difference in quinone content between various types of sludge, rather than for absolute values, which are affected by the estimation of response factors and by the matrix effect. To mitigate the latter and improve quantification accuracy, an additional step could be incorporated into the sample preparation process, such as solid-phase extraction using silica cartridges. It should be noted that different types of matrices are likely to affect distinct quinones differently. As an initial approach, the deuterated quinones used in our study provide a straightforward means of evaluating the matrix effect.

Currently, most quinones identified by our method are characterised only by their high-resolution molecular mass and distinctive fragmentation pattern, in line with previous studies [22,34]. However, this approach cannot distinguish potential isomers, so definitive annotation is not possible. Complete chemical characterisation of purified quinones is required for this. MK8:6 and MK9:5, which are prevalent in sludge samples, would be particularly interesting to study in order to determine the position of the double bonds in their side chains.

Previous studies have detected up to 26 quinones in sewage sludges and effluents [21,25,28,29]. With 57 distinct quinones detected, our method more than doubled this number and extended the range from UQ5:5-UQ11:11 and MK5:5-MK12:12 to UQ2:2-UQ12:12 and MK4:4-MK15:15, respectively. The recently characterized methyl-plastoquinones are synthesized by *Nitrospira* species [45,46], which are present in activated sludge [47]. Accordingly, we detected mPQ7:7 to mPQ10:10 (Figure 3B and Table S6), suggesting that mPQs could serve as a biomarker for nitrifying communities, which play a pivotal role in nitrogen removal during wastewater treatment [47,48]. Similarly, the dominance of UQs in sludge samples (Figure 3B) reflects the abundance of bacteria from the phylum *Pseudomonadota* [47,49].

Given that activated sludge systems are estimated to harbour up to one billion bacterial species [47], most of which are uncultured or uncharacterised, it is likely that new quinones remain to be discovered, as recently postulated [46]. Our method provides a foundation for future approaches that aim to identify novel quinones. In addition to its application in wastewater treatment [24] and microbial ecology, this method shows promise in the field of human health research, where quinone profiling could provide valuable insights into the dynamics of various human microbiomes [32,50–53].

## 5 CONCLUSION

The targeted UHPLC-Orbitrap method achieves the analysis of the most comprehensive set of isoprenoid quinones to date in just 14 minutes of chromatographic separation. Quantitative analysis was possible for 12 quinones that were available as standards, while semi-quantitative analysis of the other quinones was possible thanks to estimated response factors. Sensitivity ranged from 55.2 to 0.3 femtomoles, depending on the quinone type. When applied to sewage sludges, our method detected 57 distinct quinones and revealed different quinone profiles at various stages of wastewater treatment. This method can be used to profile quinones in complex samples and has significant potential to advance microbial ecology, environmental monitoring and biotechnological applications.

## Supporting information

Supplementary material including Text S1 and 12 supplemental figures

Table S1: PRM inclusion list

Table S2: Correspondence between wet weight and dry weight for sludge samples

Table S3: Estimated trendline coefficient for non-standard quinones

Table S4: Untargeted detection of the quinone standard mix

Table S5: Quinone detection in untargeted and targeted modes

Table S6: Quinone profiles of sludges overs weeks

## Authors’ contributions

**Morgane Roger-Margueritat**: Writing – original draft, Conceptualization, Data curation, Formal analysis, Investigation, Methodology, Validation, Visualization, **Arthur Réveillard**: Writing – original draft, Investigation, Methodology, Data curation, Formal analysis, Visualization, **Caroline Plazy**: Writing – review and editing, Formal analysis, Methodology, **Andrei Octavian Filimon**: Investigation, **Ahcène Boumendjel**: Writing – review and editing, Resources, Supervision, **Volker F. Wendisch**: Writing – review and editing, Resources, **Valérie Cunin**: Resources, **Sophie S. Abby**: Writing – review and editing, Formal analysis, Supervision, Visualization, Funding acquisition, **Audrey Le Gouellec**: Writing – review and editing, Supervision, Resources, Funding acquisition, **Fabien Pierrel**: Writing – original draft, Conceptualization, Project administration, Funding acquisition, Supervision, Investigation, Formal analysis, Validation, Visualization.

## Data availability

The quinone spectral library is available in Figshare (10.6084/m9.figshare.33924967), together with the Python script used to filter the TraceFinder output table, and the R script used for PCA and the statistical analyses. Raw MS files have been deposited in MassIVE (MSV000103362).

## Acknowledgements

The authors acknowledge Irene Krahn for her technical assistance in cultivating *C. glutamicum* Ubi5-Pd cells, and Nelle Varoquaux for her support with some computational analyses. We are also grateful to Téo Hebra for his fruitful comments on an earlier version of this manuscript, and to the staff of the wastewater treatment plant of Romans-sur-Isère, for their help with the access and the collection of sludge samples, which was performed by Olivier Réveillard. This project was funded by the French Research Agency (ANR) via the QUINEVOL project (grant agreement ANR-21-CE02-0018), as well as by the Université Grenoble Alpes via the IRGA PEE 2023 project Quinobiote. Mass spectrometry analyses were carried out at Gemeli GExiM, a clinical metabolomics Platform supported by IDEX and Auvergne Rhone Alpes Grants. Dr Audrey Le Gouellec acknowledges financial support from the Université Grenoble Alpes Foundation and from the association “Vaincre la mucoviscidose”.

## Declarations

The authors declare no competing interests. The authors confirm that the data supporting the findings of this study are available within the article and its supplementary materials.

