## Supplementary material including Text S1 and 12 supplemental figures for "Isoprenoid quinone profiling using a novel semi-quantitative HPLC-MS/MS method: application to wastewater sludges"

**Text S1: Purification of quinones from biological extracts**

**Figure S1: Quinone purification**

**Figure S2: MS/MS fragmentation spectra of quinones from the standard mixture**

**Figure S3: Quinone standard linearity using an untargeted approach**

**Figure S4: Quinones monitored with the targeted methods**

**Figure S5: PRM chromatogram of quinone standards**

**Figure S6: PRM peak integration of quinone standards with TraceFinder 4.1**

**Figure S7: Correlation between trendline coefficient and quinone chain length**

**Figure S8: Matrix effect of sludges and extraction losses estimated by the recovery of deuterated quinones**

**Figure S9: Quinone types detected in wastewater sludges**

**Figure S10: Principal Component Analysis (PCA) visualizing the quinone profiles of sludges without Hellinger transformation**

**Figure S11: Principal Component Analysis (PCA) visualizing the quinone profiles of sludges over weeks**

**Figure S12: Principal Component Analysis (PCA) visualizing the quinone profiles of primary sludges over weeks**

**Text S1**:

To further increase the diversity of standards, particularly long-chain and unsaturated MKs as well as medium-chain UQs, we purified UQ6:6 from *Saccharomyces cerevisiae* and MK10:10, MK10:9, MK11:11 and MK11:10 from *Corynebacterium glutamicum* Ubi5-Pd cells [34]. UQ6:6 was the only quinone detected in *S. cerevisiae* extracts and was separated from other lipids, including ergosterol, by silica column chromatography (Figure S1A). From 42 g of fresh yeast, purification yielded 500 μL of UQ6:6 at 1.44 mM (see methods). Extracts from *C. glutamicum* Ubi5-Pd contained comparable amounts of MK10:10, MK10:9 and MK11:10, with lower levels of MK11:11 (Figure S1B). After purification on a C18 column (see methods), we obtained 1.1 mL of MK11:10 at 0.572 mM, and 2 mL of a mixture of MK10:10 (0.162 mM) and MK10:9 (0.180 mM). Both solutions also contained small amounts of MK11:11 (0.027 mM in the MK11:10 solution and 0.011 mM in the MK10:10/MK10:9 solution). By combining these purified compounds with the commercially available standards, an equimolar mixture of 16 quinones was assembled, with MK11:11 included at sub stoichiometric levels (1.07 µM when the other quinones are at 10 µM).

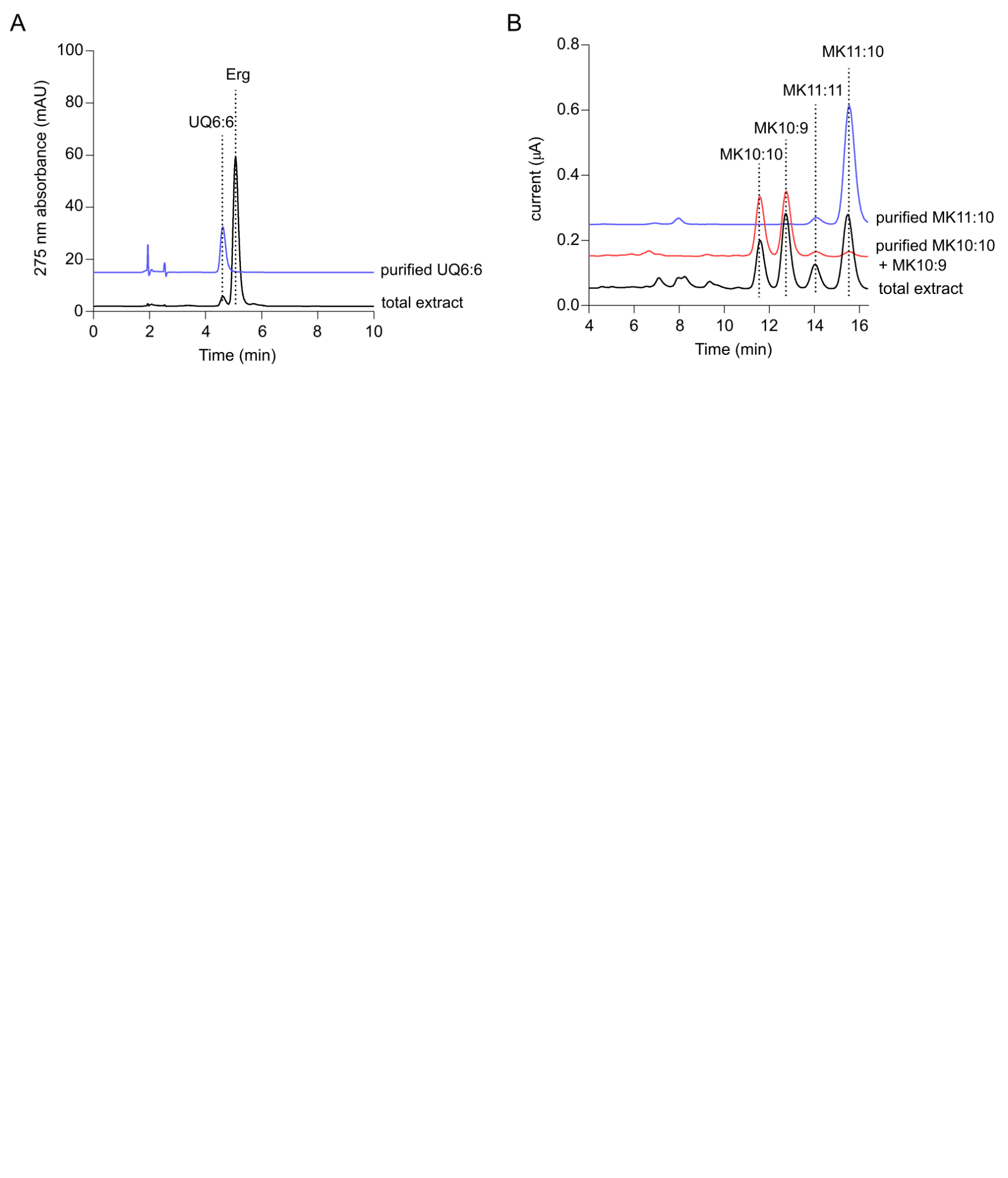

**Figure S1: Quinone purification. A.** Overlay of HPLC-UV analyses (mAU= milli absorbance units) of the total extract from *S. cerevisiae* and of the purified UQ6:6. Erg: ergosterol. **B.** Overlay of HPLC-ECD analyses of the total extract from *C. glutamicum* Ubi5-Pd cells and of the purified quinone solutions. The peaks corresponding to various quinones are marked.

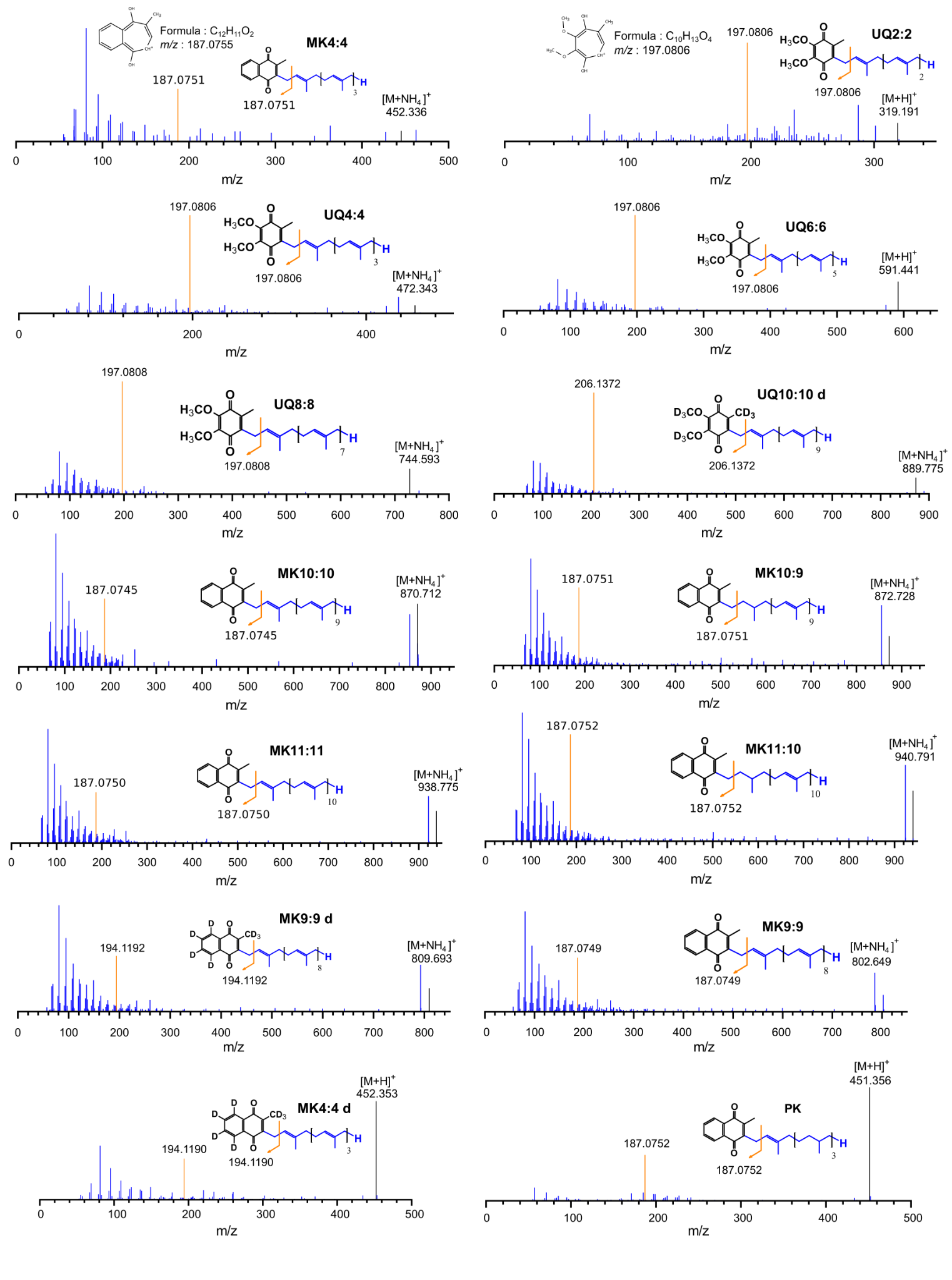

**Figure S2: MS/MS fragmentation spectra of quinones from the standard mixture.** The unfragmented quinone peak is coloured in black and the tropylium ion peak is in orange, with its structure displayed for MK and UQ. The position of the saturation on the side chains of MK11:10 and MK10:9 has not been definitively determined and is depicted arbitrarily.

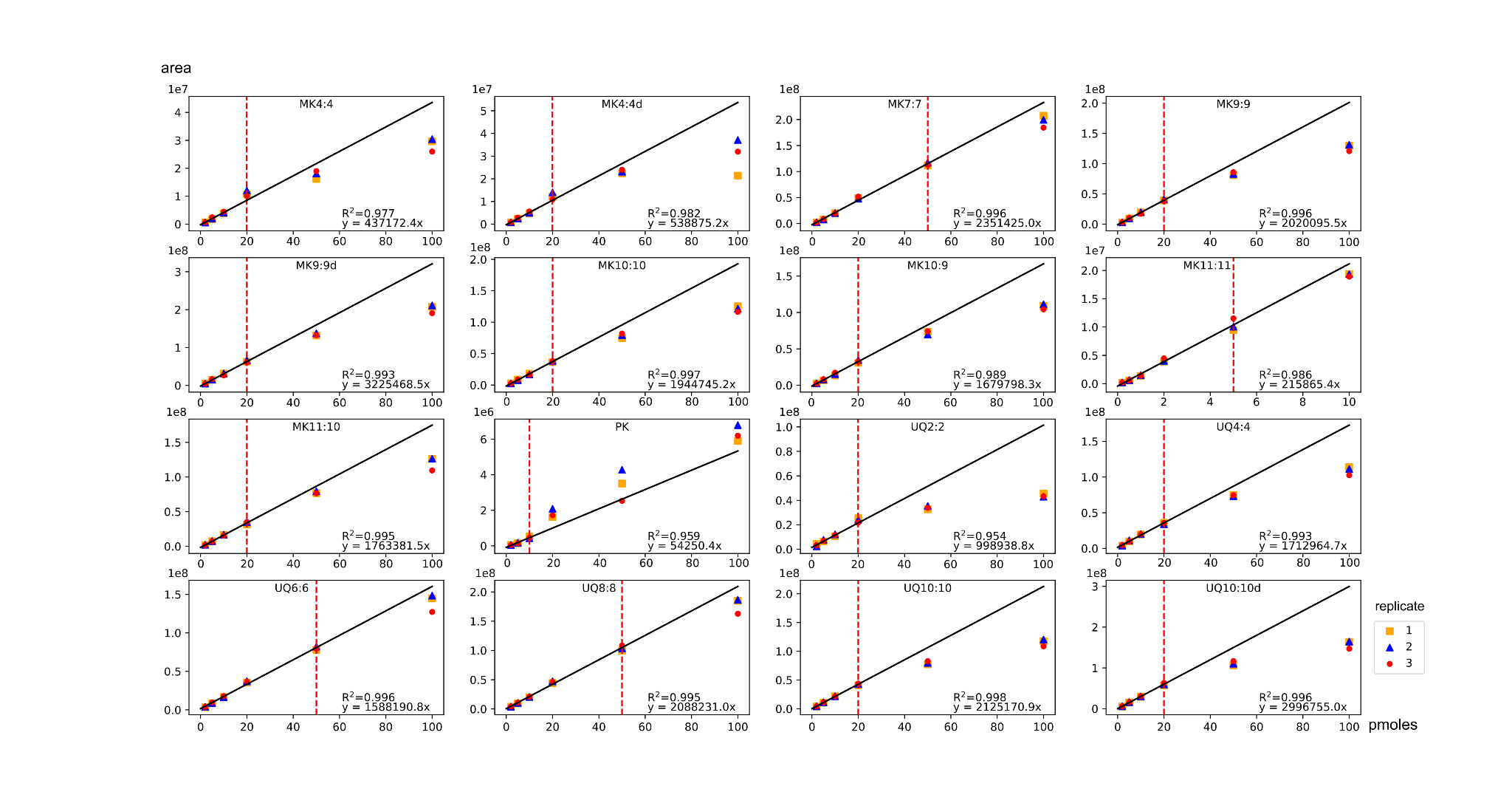

**Figure S3: Quinone standard linearity using an untargeted approach.** The quinone standard mixture was injected in a range between 0.0002 and 100 pmoles in triplicates. R² and the corresponding trendline for the linear part of the curves were computed using the Python modules numpy. The red dashed line corresponds to the Upper Limit of Quantification (ULOQ).

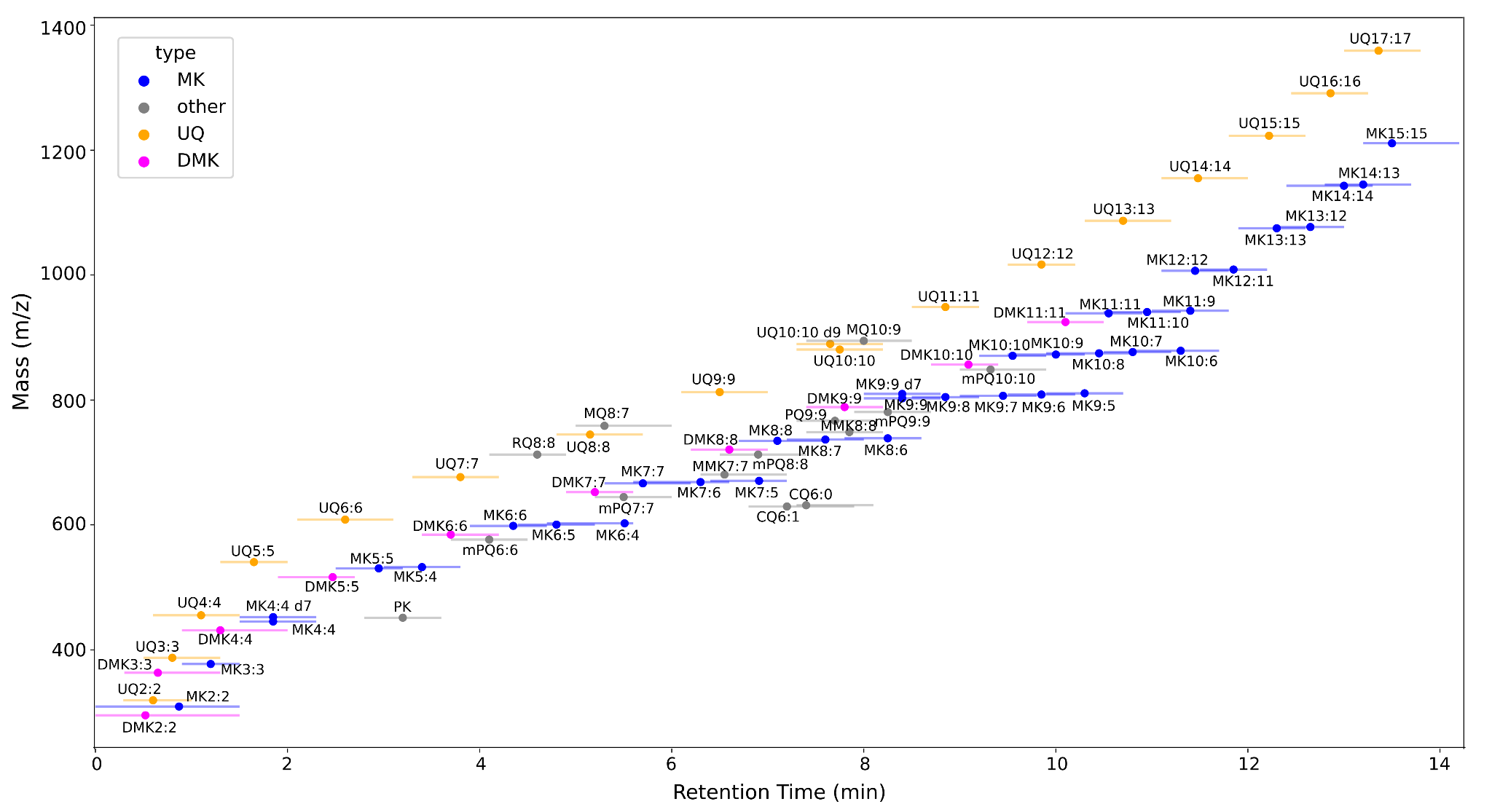

**Figure S4: Quinones monitored with the targeted method.** Each compound mass (m/z) with their corresponding windows (min) were extracted based on the PRM inclusion list (Table S1) and the plot was obtained with the python module matplotlib. UQ: ubiquinone; DMK: demethylmenaquinone; MK: menaquinone; PK: phylloquinone; RQ: rhodoquinone; mPQ: methyl- plastoquinone; PQ: plastoquinone; CQ: caldariellaquinone; MQ: methylene ubiquinone; MMK: methyl-menaquinone.

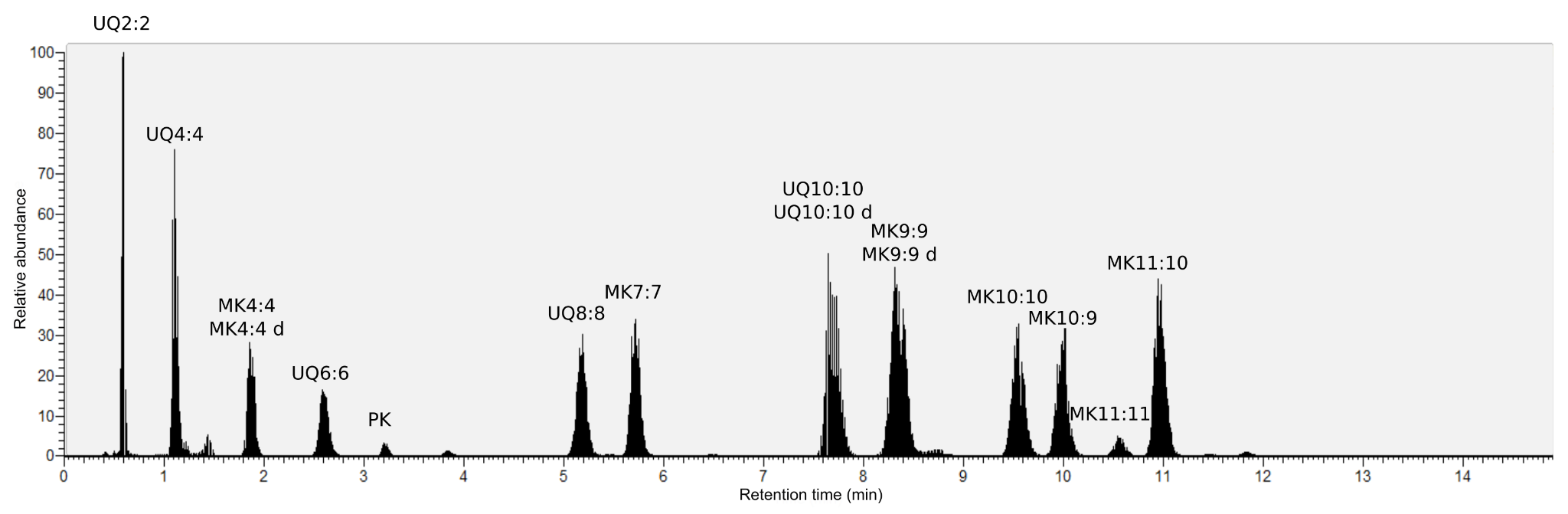

**Figure S5: PRM chromatogram of quinone standards.** 20 pmoles of the quinone standard mixture were injected and the raw data was visualized with the Qual Browser option of Xcalibur (Thermo Fisher).

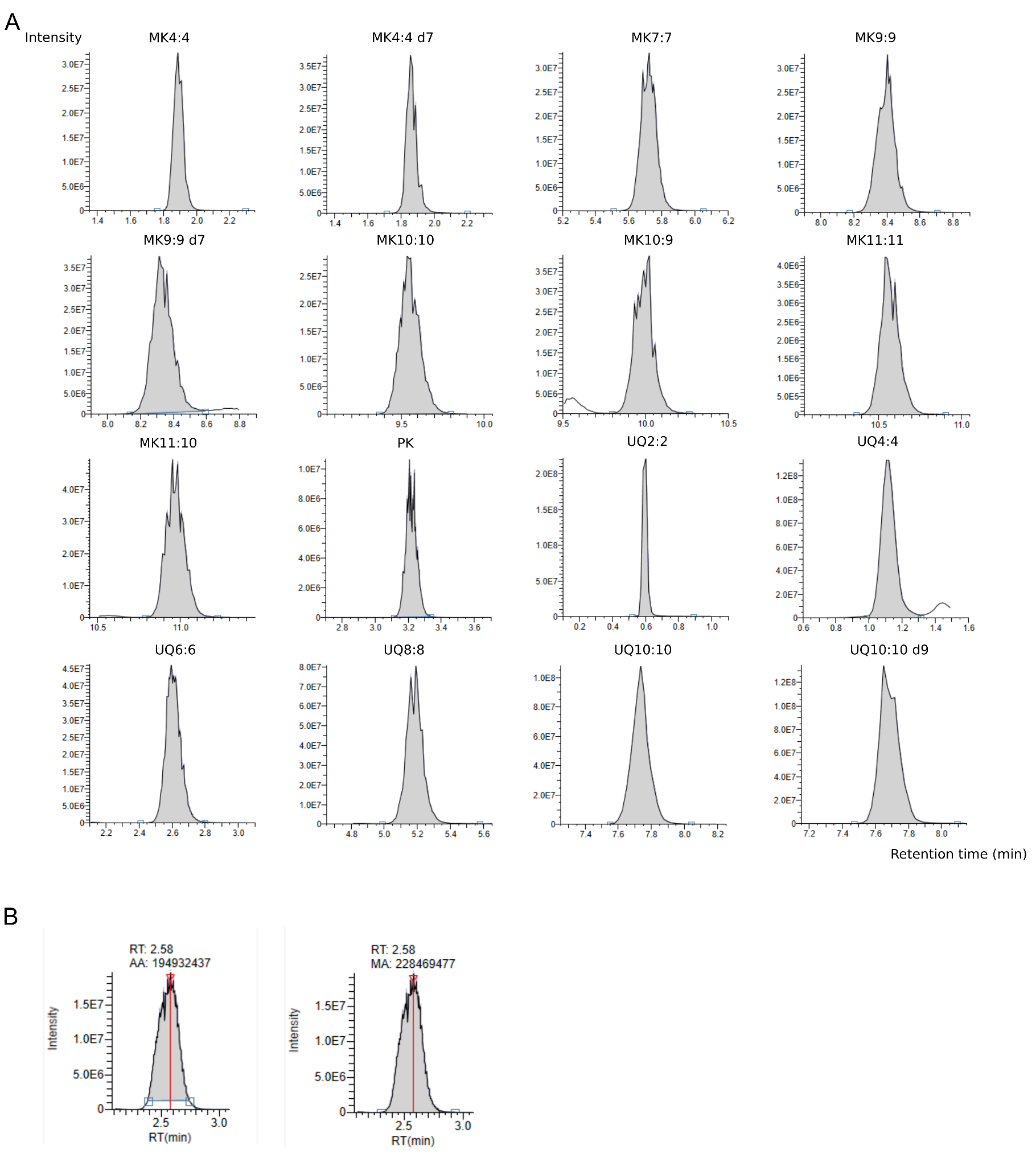

**Figure S6: PRM peak integration of quinone standards with TraceFinder 4.1.** **A**. 20 pmoles of the standard mixture were injected and the raw data were analyzed with a Master method designed with TraceFinder 4. **B**. When necessary, manual curation was carried out to integrate the entire peak when it had not been fully recovered automatically as it is shown with a peak before (left) and after (right) the curation.

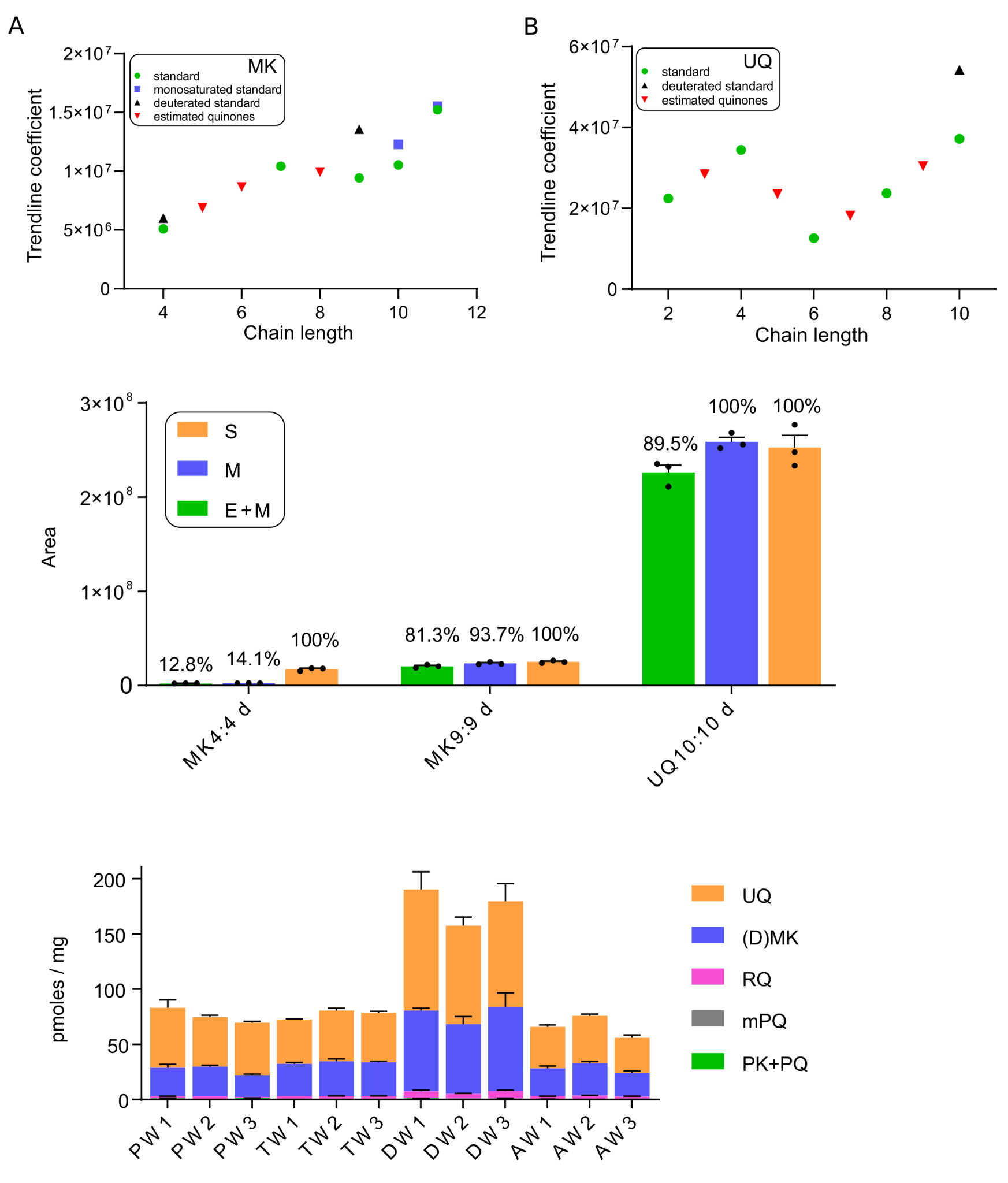

**Figure S7: Correlation between trendline coefficient and quinone chain length.** The trendline coefficient of each standard quinone (see Table 1) was plotted against the chain length for either menaquinone (**A**) or ubiquinone (**B**). The linear regression was computed and the trendline and correlation coefficients (R²) are displayed. MK: menaquinone; UQ: ubiquinone.

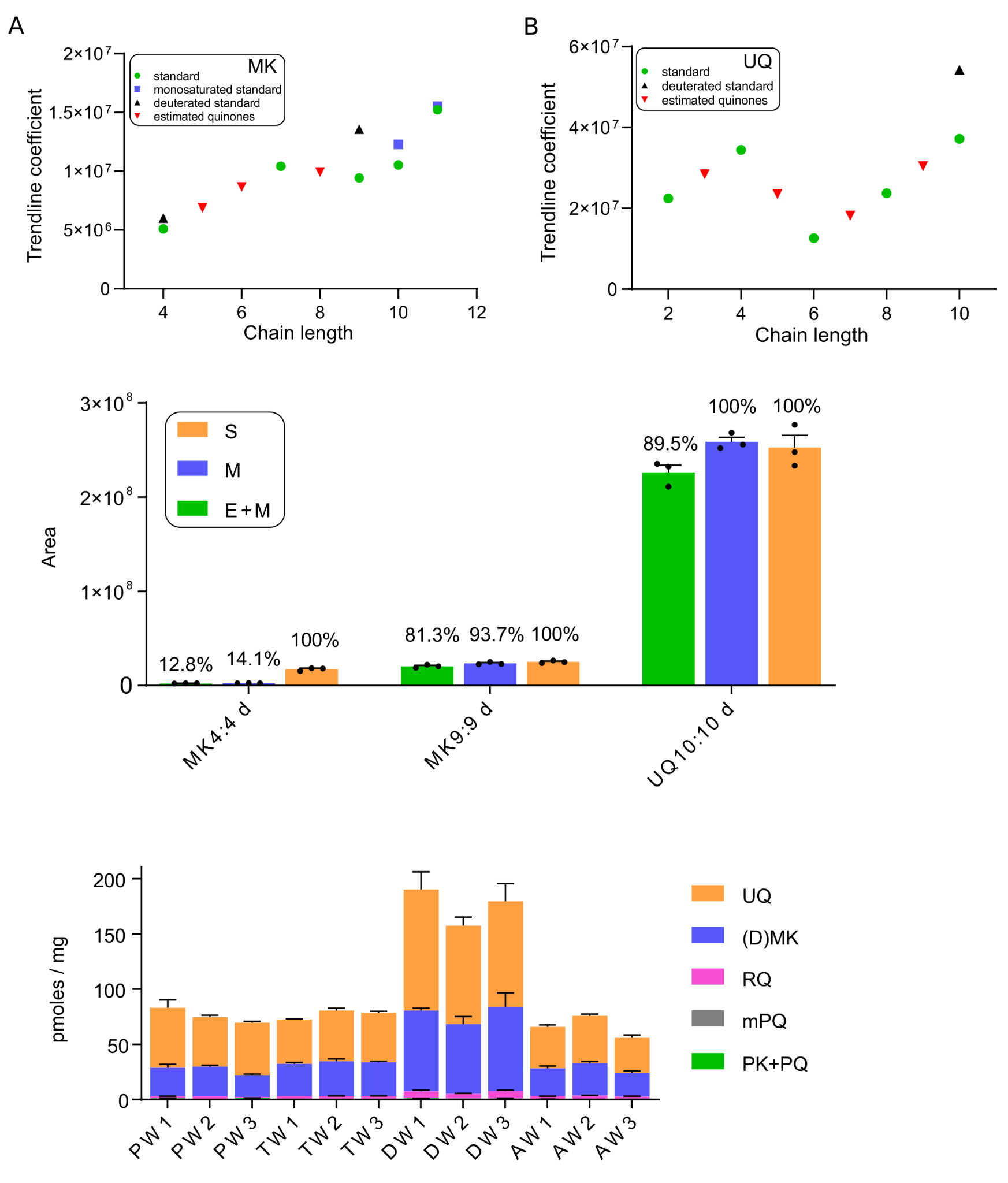

**Figure S8: Matrix effect of sludges and extraction losses estimated by the recovery of deuterated quinones.** A mix of the 4 sludge types (primary, thickened, activated or dehydrated) from week 1 was pooled for a neutral lipid extraction step. Deuterated quinones were added either before (green) or after (blue) the extraction to assess extraction losses and matrix effect, respectively. The peak area obtained after HPLC/MS-MS analysis was compared to that obtained after analysis of the pure deuterated mixture (orange). The percentage indicates the recovery of each deuterated quinone. Mean ± SD (n = 3). S: Standard; M: Matrix effect; E + M: Matrix effect + Extraction losses.

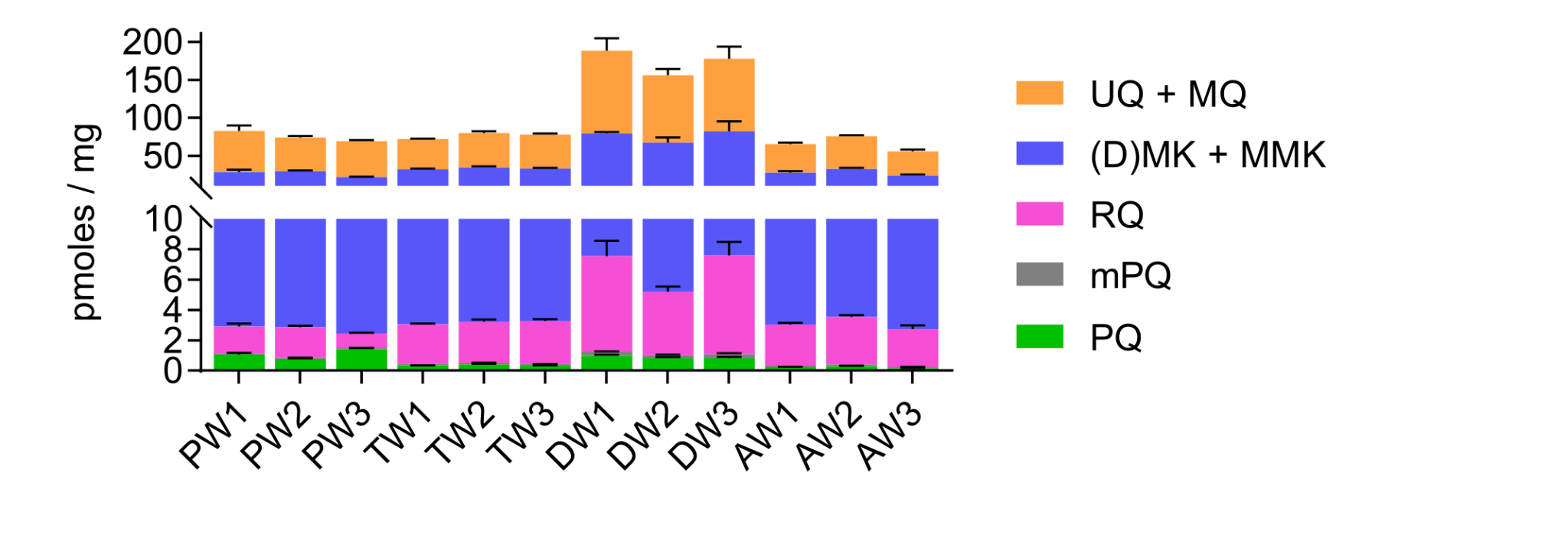

**Figure S9: Quinone types detected in wastewater sludges.** Quinones from the primary (P), thickened (T), activated (A) or dehydrated (D) sludges retrieved over 3 weeks (W1, 2 or 3) were quantified (pmoles/mg of sludge wet weight) and grouped according to their types: ubiquinone (UQ) + methylene ubiquinone (MQ), (demethyl)menaquinone (D)MK + methylmenaquinone (MMK), rhodoquinone (RQ), methyl-plastoquinone (mPQ) and phylloquinone (PK) + plastoquinone (PQ). Mean 土 SEM (n = 2-6).

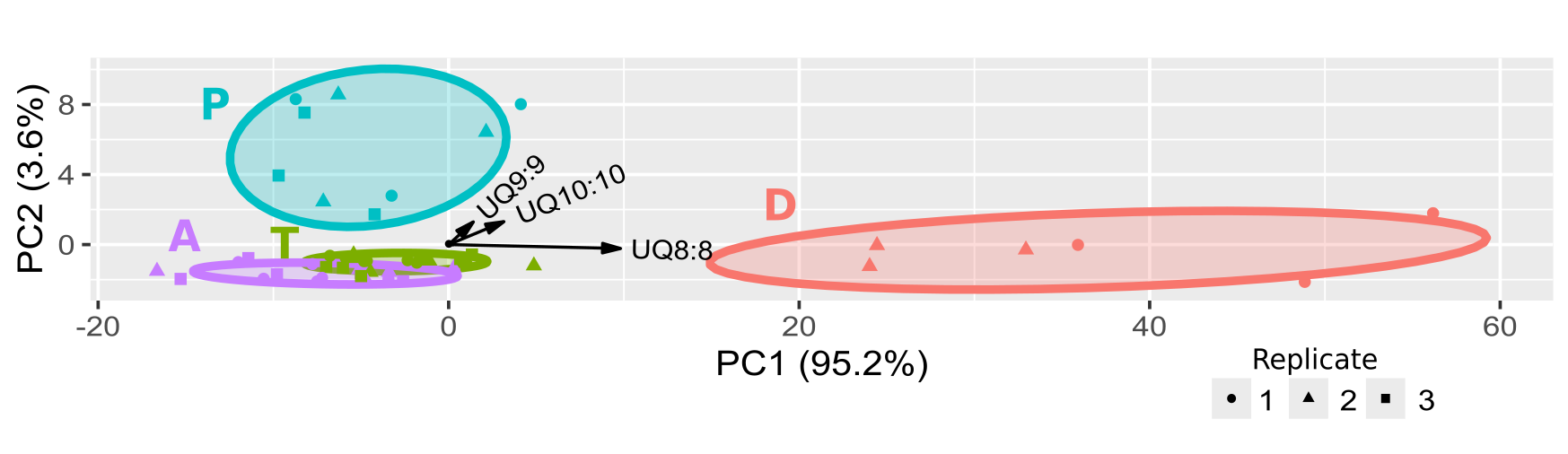

**Figure S10: Principal Component Analysis (PCA) visualizing the quinone profiles of sludges without Hellinger transformation.** Sludge samples collected in four treatment steps (primary (P), thickened (T), activated (A) and dehydrated (D) sludges) were analysed and quinones were quantified. PCA and the circles were computed and visualized with the ggbiplot module of R with the best quinone descriptors displayed by black arrow.

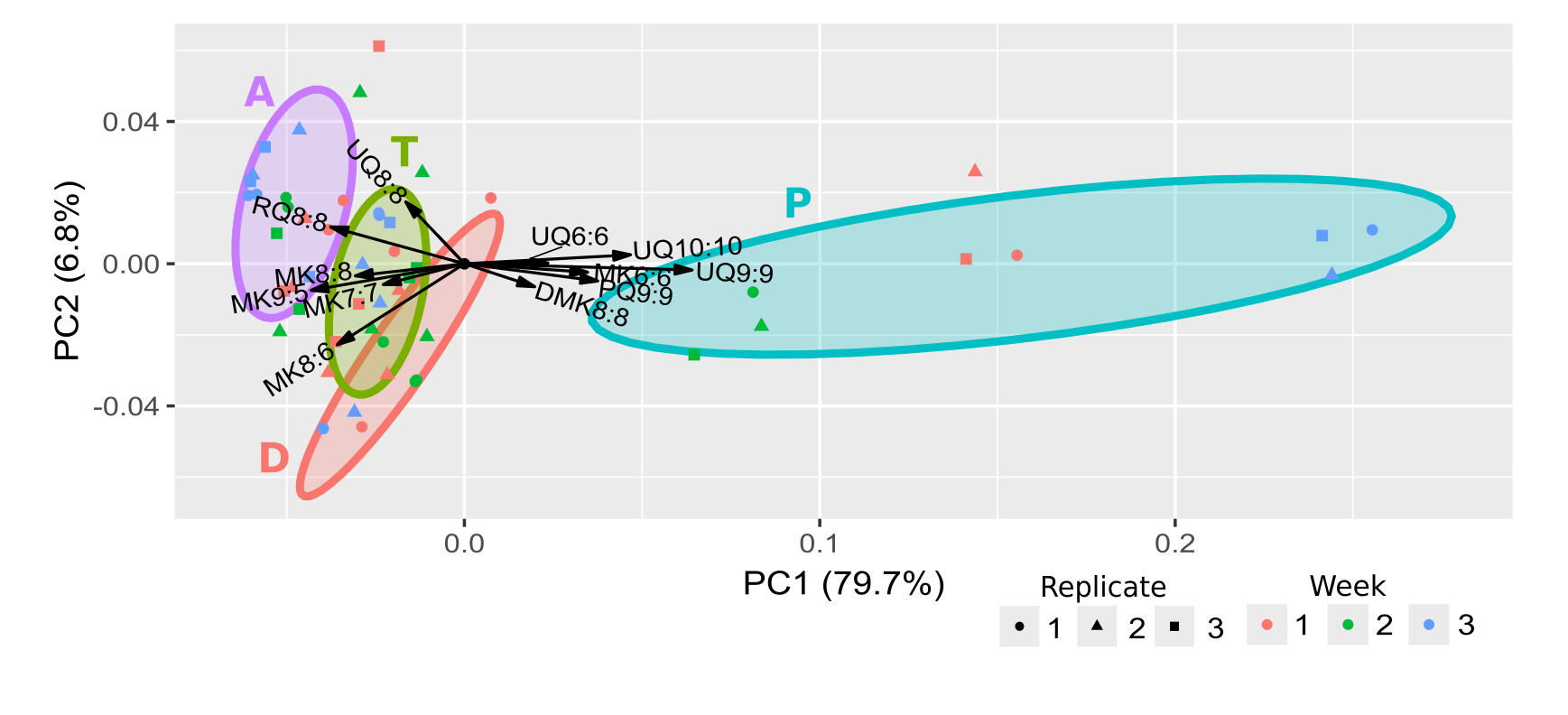

**Figure S11: Principal Component Analysis (PCA) visualizing the quinone profiles of sludges over weeks.** Sludge samples collected during three weeks (respectively in orange, green and blue) in each sludge type (primary (P), thickened (T), activated (A) and dehydrated (D)) were analysed and quinones were quantified. Hellinger transformation was applied on the values and PCA was computed and visualized with the ggbiplot module of R with the best quinone descriptors displayed by black arrow.

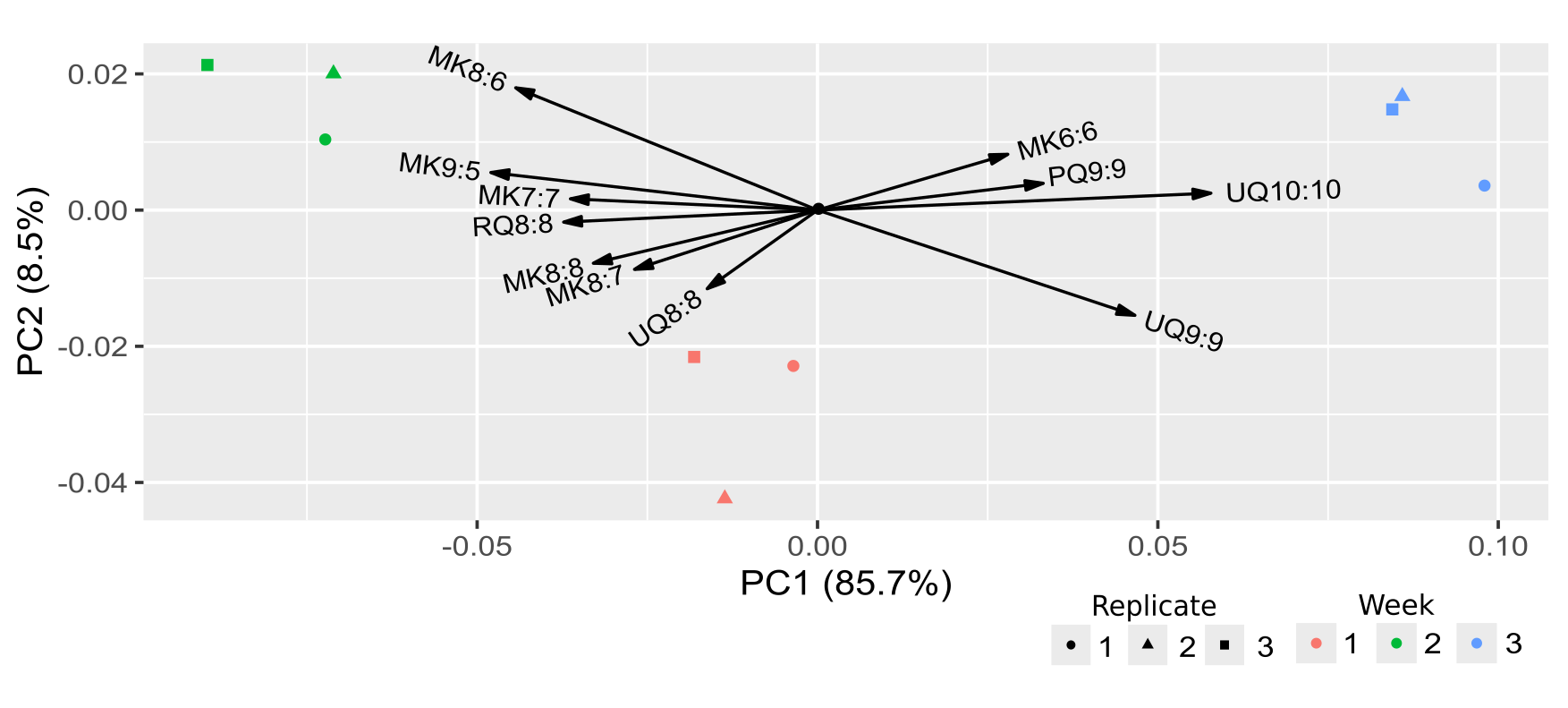

**Figure S12: Principal Component Analysis (PCA) visualizing the quinone profiles of primary sludges over weeks.** Primary sludge samples collected during three weeks (respectively in orange, green and blue) were analysed and quinones were quantified. Hellinger transformation was applied on the values and PCA was computed and visualized with the ggbiplot module of R with the best quinone descriptors displayed by black arrow.
